# Genome reduction is associated with bacterial pathogenicity across different scales of temporal and ecological divergence

**DOI:** 10.1101/2020.07.03.186684

**Authors:** Gemma G. R. Murray, Jane Charlesworth, Eric L. Miller, Michael J. Casey, Catrin T. Lloyd, Marcelo Gottschalk, A. W. (Dan) Tucker, John J. Welch, Lucy A. Weinert

**Affiliations:** Department of Veterinary Medicine, University of Cambridge, CB3 0ES, UK; Department of Genetics, University of Cambridge, CB2 3EH, UK; Warwick Medical School, University of Warwick, CV4 7AL, UK; Haverford College, Pennsylvania, 19041, USA; School of Mathematical Sciences, University of Southampton, SO17 1BJ, UK; Département de pathologie et microbiologie, Université de Montréal, 3200, Canada

## Abstract

Emerging bacterial pathogens threaten global health and food security, and so it is important to ask whether these transitions to pathogenicity have any common features. We present a systematic study of the claim that pathogenicity is associated with genome reduction and gene loss. We compare broad-scale patterns across all bacteria, with detailed analyses of *Streptococcus suis,* a zoonotic pathogen of pigs, which has undergone multiple transitions between disease and carriage forms. We find that pathogenicity is consistently associated with reduced genome size across three scales of divergence (between species within genera, and between and within genetic clusters of *S. suis*). While genome reduction is most often associated with bacterial endosymbionts, other correlates of symbiosis (reduced metabolic capacity, GC content, and the expansion of non-coding elements) are not found consistently in pathogens, and genome reduction in pathogens cannot be attributed to changes in intracellularity or host restriction. Together, our results indicate that genome reduction is a predictive marker of pathogenicity in bacteria, and that the causes and consequences of genome reduction in pathogens are sometimes distinct from those in endosymbionts.

## 1. Introduction

The emergence of new bacterial pathogens is a major threat to human health and food security across the globe (1). Identifying common features could help us to understand, predict, and ultimately prevent these transitions to pathogenicity. One intriguing observation is that some of the most serious human pathogens have smaller genomes and fewer genes than their closest non-pathogenic or less pathogenic relatives (2–10). Nevertheless, without formal comparative studies, it is difficult to know whether these are isolated instances of genome reduction, or part of a broader trend (7, 11).

There are also doubts about whether genome reduction has anything to do with pathogenicity *per se* (7). Most notably, similar patterns of genome reduction are found in mutualist or commensal bacteria that have adopted a host-restricted or intracellular lifestyle. In these bacteria, genome reduction appears as one part of a syndrome that also includes a decreased proportion of G/C relative to A/T bases, a proportional expansion of non-coding regions (including pseudogenes and other non-functional elements), and a loss of genes in metabolic pathways. This “endosymbiont syndrome” is a plausible outcome of long-term evolutionary processes associated with small isolated populations, and greater dependency on the host (4, 12–15). As such, the genome reduction observed in some pathogens may be a reflection of their host-restricted lifestyle, rather than their pathogenicity.

Here, we present a systematic study of genome reduction and pathogenicity in bacteria, across multiple scales of divergence (Figure 1). First, at the broadest scale, we compare pairs of bacterial species, where a known vertebrate pathogen has a non-pathogenic relative in the same genus. These data span 6 distinct phyla, and a wide range of ecologies (Figure 1a, Table 1). Second, we focus on *Streptococcus suis*, an opportunistic pathogen of pigs, whose pathogenicity has previously been associated with genome reduction (5, 16, 17). This bacterium is of particular interest because it has undergone multiple independent transitions between carriage and disease forms, yet both forms are generally extracellular, and have equal levels of host restriction. As such, we can observe “replicated” changes in pathogenicity, that are not accompanied by changes in the broader ecology (see below). Furthermore, the species includes multiple genetic clusters of closely-related isolates. By comparing patterns between clusters (Figure 1b), and between isolates within clusters (Figure 1c), we can contrast longer-term processes to smaller-scale or rapid genomic changes.

**Figure 1.**
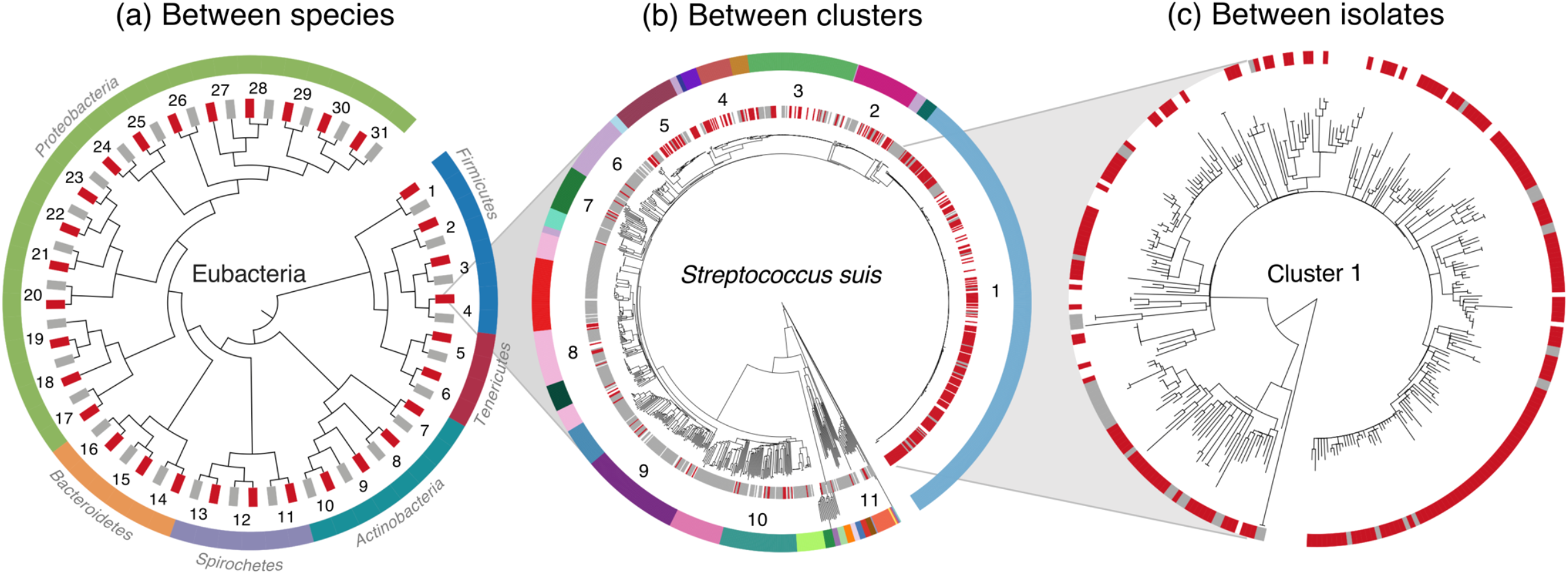
The evolution of pathogenicity over three evolutionary scales: (a) between species of bacteria, (b) between clusters of *Streptococcus suis,* and (c) between isolates of *S. suis* within clusters. (a) A cladogram of our 31 pairs of congeneric species, comprising a pathogen (red) and a non-pathogen (grey). Numbers refer to Table 1 and suprageneric relationships are from (52). (b) A core genome phylogeny of our 1,079 isolates of *S. suis.* Individual disease (red) and carriage (grey) isolates are indicated in the inner strip. The outer strip describes the 34 genetic clusters, with the 11 “mixed clusters” that include multiple disease and carriage isolates numbered. (c) An illustrative phylogeny of our largest and most pathogenic cluster, constructed from a recombination-stripped local core genome alignment. Individual disease (red) and carriage (grey) isolates are again indicated on the strip.

**Table 1.**
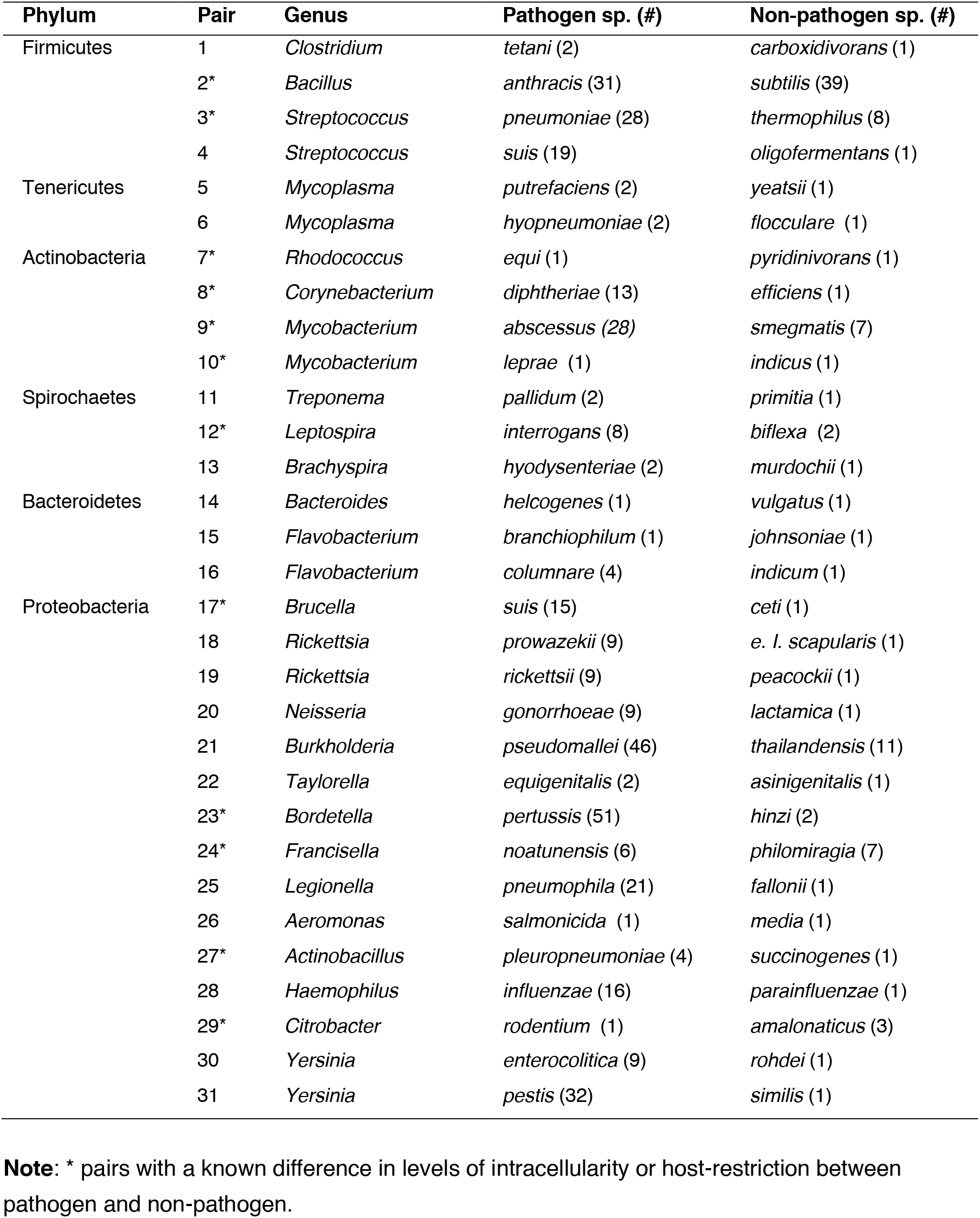
Between-species data set.

| Phylum | Pair | Genus | Pathogen sp. (#) | Non-pathogen sp. (#) |
| --- | --- | --- | --- | --- |
| Firmicutes | 1 | *Clostridium* | *tetani* (2) | *carboxidivorans* (1) |
|  | 2* | *Bacillus* | *anthracis* (31) | *subtilis* (39) |
|  | 3* | *Streptococcus* | *pneumoniae* (28) | *thermophilus* (8) |
|  | 4 | *Streptococcus* | *suis* (19) | *oligofermentans* (1) |
| Tenericutes | 5 | *Mycoplasma* | *putrefaciens* (2) | *yeatsii* (1) |
|  | 6 | *Mycoplasma* | *hyopneumoniae* (2) | *flocculare* (1) |
| Actinobacteria | 7* | *Rhodococcus* | *equi* (1) | *pyridinivorans* (1) |
|  | 8* | *Corynebacterium* | *diphtheriae* (13) | *efficiens* (1) |
|  | 9* | *Mycobacterium* | *abscessus* (28) | *smegmatis* (7) |
|  | 10* | *Mycobacterium* | *leprae* (1) | *indicus* (1) |
| Spirochaetes | 11 | *Treponema* | *pallidum* (2) | *primitia* (1) |
|  | 12* | *Leptospira* | *interrogans* (8) | *biflexa* (2) |
|  | 13 | *Brachyspira* | *hyodysenteriae* (2) | *murdochii* (1) |
| Bacteroidetes | 14 | *Bacteroides* | *helcogenes* (1) | *vulgatus* (1) |
|  | 15 | *Flavobacterium* | *branchiophilum* (1) | *johnsoniae* (1) |
|  | 16 | *Flavobacterium* | *columnare* (4) | *indicum* (1) |
| Proteobacteria | 17* | *Brucella* | *suis* (15) | *ceti* (1) |
|  | 18 | *Rickettsia* | *prowazekii* (9) | *e. l. scapularis* (1) |
|  | 19 | *Rickettsia* | *rickettsii* (9) | *peacockii* (1) |
|  | 20 | *Neisseria* | *gonorrhoeae* (9) | *lactamica* (1) |
|  | 21 | *Burkholderia* | *pseudomallei* (46) | *thailandensis* (11) |
|  | 22 | *Taylorella* | *equigenitalis* (2) | *asinigenitalis* (1) |
|  | 23* | *Bordetella* | *pertussis* (51) | *hinzi* (2) |
|  | 24* | *Francisella* | *noatunensis* (6) | *philomiragia* (7) |
|  | 25 | *Legionella* | *pneumophila* (21) | *fallonii* (1) |
|  | 26 | *Aeromonas* | *salmonicida* (1) | *media* (1) |
|  | 27* | *Actinobacillus* | *pleuropneumoniae* (4) | *succinogenes* (1) |
|  | 28 | *Haemophilus* | *influenzae* (16) | *parainfluenzae* (1) |
|  | 29* | *Citrobacter* | *rodentium* (1) | *amalonaticus* (3) |
|  | 30 | *Yersinia* | *enterocolitica* (9) | *rohdei* (1) |
|  | 31 | *Yersinia* | *pestis* (32) | *similis* (1) |

**Note:** \* pairs with a known difference in levels of intracellularity or host-restriction between pathogen and non-pathogen.

## 2. Results

### Data sets

For our broadest-scale between-species data set (Figure 1a), we carried out a systematic search for phylogenetically-independent species pairs, comprising a pathogen, a congeneric non-pathogen, and an outgroup, with publicly available whole-genome data (see methods for details). Our criteria left us with 31 species pairs (Tables 1, S1) represented by 478 ingroup genomes (Table S2). Because these pairs differed both in their sampling densities (i.e. the number of available genomes), and in the evolutionary distances between the species, we developed a phylogenetic comparative method to correct for these differences.

For our within-species comparisons (Figure 1b-c), we collated 1,079 whole genomes of *S. suis,* collected across three continents (Tables 2, S3). Data were associated with clinical information, and include “carriage isolates” from the tonsils of pigs without *S. suis* associated disease, and “disease isolates” from the site of infection in pigs with *S. suis* associated disease (divided into respiratory or systemic infections). We also included zoonotic disease isolates from humans with systemic disease in Vietnam, which a previous study found to be indistinguishable from isolates associated with systemic disease in pigs (5). A core genome alignment was used to infer a consensus phylogeny; but as *S. suis* is highly recombining, we inferred genetic clusters using an approach that is agnostic of the phylogeny (18, 19). We identified 34 clusters, of which 33 contained isolates that could be unambiguously categorised as carriage or disease (Figure 1b, Figure S1a, Table S4). These clusters have variable levels of genetic diversity and include some with recent origins; for example, a previous study dated the origin of our largest and most pathogenic cluster (cluster 1, Figure 1c) to the 1920s (5). There were marked differences between clusters in the proportion of disease isolates, and this correlated with the presence of virulence genes and serotypes with known disease associations (Figure S2). Allowing for comparisons at the finest scale, 11/34 were “mixed clusters”, containing both multiple carriage isolates and multiple disease isolates, and these are numbered in Figure 1b.

**Table 2.** *Streptococcus suis* data set.

| Origin | # Isolates:<br>Total (SP, RP, C) | Genome size<br>range (Mb) | Cluster(s) | Date(s) | Reference(s) |
| --- | --- | --- | --- | --- | --- |
| UK | 440 (43, 52, 205) | 1.95 - 2.56 | 1-11, 13-14, 16-18, 20,<br>22-25, 27-28, 30-33 | 2009 - 2015 | (5, 31) |
| Canada | 197 (36, 31, 56) | 1.92 - 2.49 | 1-3, 5-18, 20, 22-26,<br>29-30 | 1983 - 2016 | (33) |
| China | 197 (0, 0, 197) | 2.06 - 2.54 | 1, 3, 5-12, 18-22, 27-30,<br>32, 34 | 2013 - 2014 | (31) |
| Vietnam | 190 (149, 0, 32) | 1.97 - 2.19 | 1 | 2000 - 2010 | (5) |
| USA | 16 (3, 4, 0) | 2.02 - 2.46 | 3, 5, 8-10, 26 | 2016 | (33) |
| Spain | 10 (7, 0, 0) | 2.03 - 2.42 | 1, 4, 8, 22 | 2016 | (33) |
| Reference<br>collection | 29 (14, 2, 3) | 1.98 - 2.30 | 1-4, 6, 8, 13-14, 18-19 | - | (34) |

**Note:** SP: Systemic pathogen, RP: Respiratory pathogen, C: Carriage.

### Pathogenicity is associated with genome reduction at all divergence scales

Across all three scales of divergence our data sets showed substantial variation in genome size, and in all cases we found a consistent association with pathogenicity (Figure 2). At the broadest scale, pathogenic species had smaller genomes than their non-pathogenic relatives more often than expected by chance (Figure 2a; see also Table S5 and Figure S3a for robustness analyses). Within *S. suis,* genetic clusters had smaller genomes when they contained a higher proportion of disease isolates (Figure 2b; see also Table S6), and in 11/11 mixed clusters, disease isolates had smaller genomes on average, than carriage isolates from the same cluster (Figure 2c).

**Figure 2.**
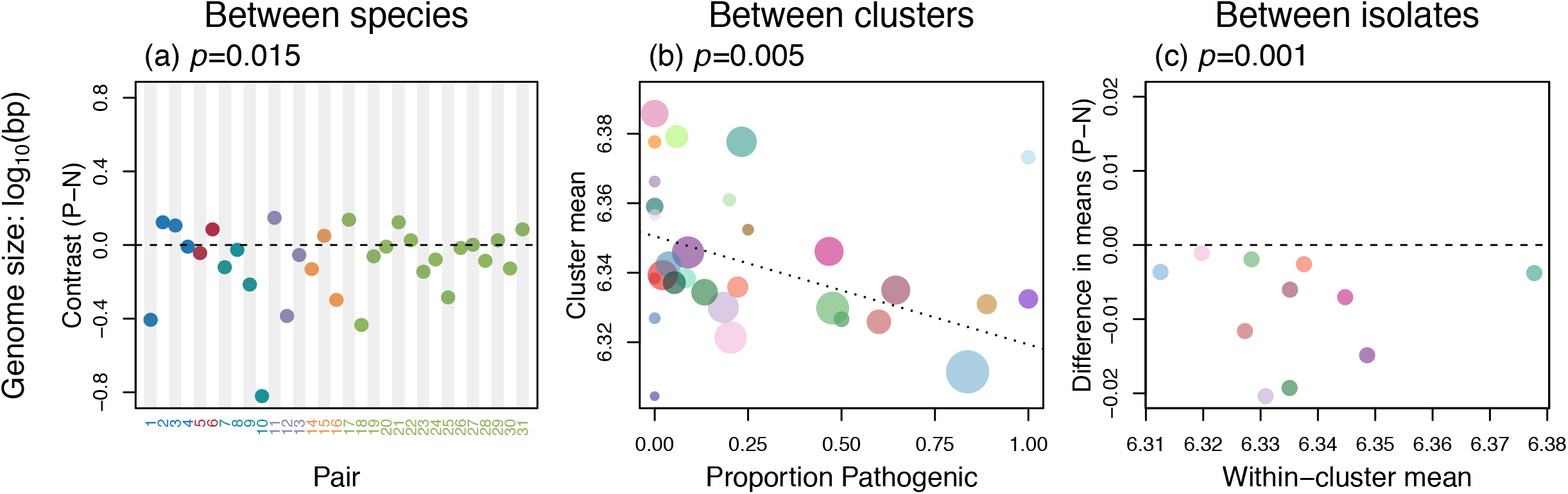
Pathogenicity is associated with smaller genomes. (a) Each point represents a phylogenetically independent species pair, with numbers and colours from Figure 1a. Standardised contrast values are ordered such that negative values imply smaller genomes in the pathogenic species. *p*-value is from a permutation test of the null hypothesis of no difference in the mean contrast value (as indicated by the dotted line). (b) Each point represents a cluster of *S. suis* isolates, with colours from Figure 1 b, and sizes indicating the number of isolates. A sample-size weighted regression shows that clusters containing a larger proportion of pathogenic isolates have a smaller average genome size. (c) Each point shows the difference in mean genome size between disease and carriage isolates in a cluster of *S. suis* isolates, such that negative values imply smaller genomes in the pathogens. Each point corresponds to a “mixed cluster” (containing multiple disease and carriage isolates), with colours and numbers matching Figure 1b. The *p*-value is from a permutation test as in (a).

Three further lines of evidence suggest that changes in genome size persist over long periods of time. First, in our between-species data set, a Brownian model of genome size evolution provides a good fit, such that longer times lead to larger changes (Figure S4). Second, in *S. suis*, between-cluster differences in genome size remain apparent when we consider the carriage isolates alone: carriage isolates have smaller genomes when they are found in clusters containing a higher proportion of disease isolates (“Dataset C” in Table S6). Finally, in *S. suis,* there is an association between genome size, and disease severity. We see the smallest genomes in isolates associated with more invasive systemic disease, with less severe respiratory disease isolates tending to have intermediate genome size (Figure S5).

All of these results apply to total genome size. However, we observed the same pattern when we used genome annotations to consider only functional elements. Across all divergence scales, pathogenicity was associated with fewer genes, and smaller coding length, as well as smaller genome size (Figure S3a-i).

### Pathogenicity is not consistently associated with endosymbiont syndrome

In endosymbionts, genome reduction is frequently associated with a preferential loss of metabolic genes, proliferation of non-functional DNA, and low GC content. In our data, evidence of this “endosymbiont syndrome” is patchy.

Over all three timescales, pathogens do contain fewer genes with metabolic functions (Figure S3m-o; Table S5), but not more so than expected, given the overall pattern of gene loss. In particular, the proportion of genes that have a metabolic function does not differ significantly between pathogenic and non-pathogenic congeners (Figure 3a, Table S5). While in *S. suis,* the proportion of metabolic genes is actually higher in pathogens (Figure 3b-c) – the opposite of the predicted pattern.

**Figure 3.**
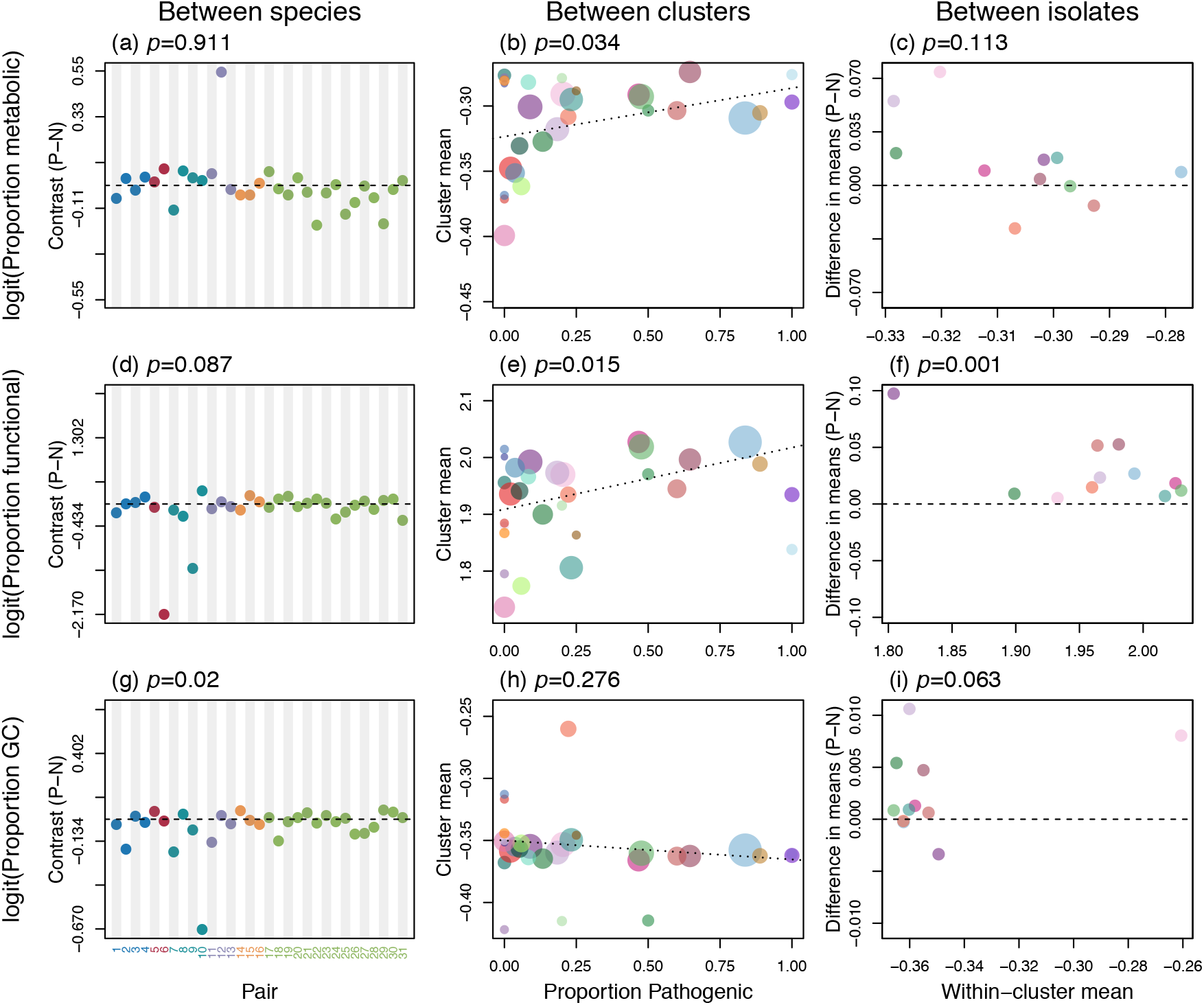
Pathogenicity is not consistently associated with the “endosymbiont syndrome”. Each panel tests for an association between pathogenicity and another signature of the endosymbiont syndrome, (a-c): the proportion of protein-coding genes with metabolic function, (d-f): the proportion of the genome with known function, (g-i): the proportion of the genome comprising GC base pairs. Negative contrasts (a,c,d,f,g,i) or negative regression slopes (b,e,h) are consistent with an endosymbiont syndrome. All other details match Figure 2.

Results for non-coding regions are similar. Between species, there is no consistent difference between pathogens and non-pathogens in the proportion of the genome that is functional (Figure 3d, Table S5, Figure S6). While in *S. suis,* and against predictions, the functional proportion is higher in the disease isolates (Figure 3e-f).

For GC content, the third component of endosymbiont syndrome, results are more complicated. Between species, we did observe the predicted tendency for pathogens to be GC-poor (Figure 3g). However, further analysis showed that this result was attributable to a subset of pairs that had undergone a large-scale shift in ecology. In 12/31 pairs, the pathogenic species showed a greater degree of host restriction and/or intracellularity than its non-pathogenic congener (Table 1), and the relationship between pathogenicity and GC held only for these pairs, and not for the remainder of the data (Table S5). Results for genome size showed the exact opposite pattern: the association between genome reduction and pathogenicity was driven by 19/31 pairs without a clear difference in these aspects of ecology (Table S5).

Within *S. suis,* the raw results for GC showed no clear patterns (Figures 3h,i, S1c). However, this is attributable to divergent groups of S. suis with unusual GC content (20; Figures S1a,c, S7a). Once these groups are removed, results suggest a balance of opposing forces. Smaller *S. suis* genomes tend to rapidly lose GC-poor mobile elements, while slowly accumulating GC-to-AT base substitutions in their core genomes (see Figure S7 for full details). The result is that the predicted association between pathogenicity and low GC is observed, but only between clusters, and only in the core genome (Figure S8).

### The link between pathogenicity and genome reduction in *Streptococcus suis*

Genome reduction might be a cause of pathogenicity or a consequence (7). In *S. suis,* where we can examine the process over a range of timescales, the evidence is ambiguous.

On the one hand, we have shown that genome reduction in *S. suis* can occur rapidly and without other signatures of the endosymbiont syndrome (Figures 2c, 3c,f,i). This is consistent with genome reduction having a causal role in pathogenicity. However, we can rule out one causal hypothesis. In other bacteria, the loss of immune targets or other “anti-virulence” genes have been linked to increased pathogenicity (21). Despite the overall trend for gene loss, our *S. suis* data show no evidence that particular genes are preferentially absent in disease isolates or pathogenic clusters (Figure S10). Instead, we find that genes are preferentially present in more pathogenic clusters – with putative virulence factors bucking the overall trend (Figures S2, S10; 5).

We have also shown that genome reduction in *S. suis* persists for long periods (Figure 2b), consistent with it being an ongoing consequence of pathogenicity. This would require some difference between the ecology of pathogens and non-pathogens, despite there being no difference in host restriction or intracellularity. One possibility is that pathogens have higher rates of transmission. There is evidence for this in the very broad geographic spread of the more pathogenic clusters of *S. suis,* despite their relatively recent origin. For example, cluster 1 includes isolates from China, Vietnam, the UK, Spain and Canada (Table S4). Higher transmission rates would lead to more population bottlenecks and increased levels of genetic drift (22, 23). Consistent with this hypothesis, we have already demonstrated a slow increase in the AT content of the *S. suis* core genome (Figures S7, S8); and there is further evidence of increased drift in more pathogenic clusters, namely shorter terminal branches and faster rates of protein evolution (Figure S1, Figure S9). When coupled with a mutational bias towards deletions, this could lead to genome reduction as a gradual passive consequence of pathogenicity.

## 3. Summary

We have demonstrated a statistical association between pathogenicity and genome reduction in bacteria, which applies both across bacterial phyla and across different scales of divergence. This suggests that genome reduction could prove a useful marker of emerging and increasing pathogenicity in bacteria. We have further demonstrated that genome reduction in pathogens is independent of both changes in host restriction or intracellularity, and of other signatures of genome evolution commonly observed in endosymbionts (although these signatures are not entirely absent). The consistent trend for genome reduction across different pathogenic ecologies and different scales of divergence, combined with the variable presence of other signatures of genome evolution observed in endosymbionts, could reflect no single underlying cause for genome reduction in pathogens and instead suggests that multiple features of pathogenic ecologies underpin this process.

## 4. Methods

### Datasets

For the between-species data set, we aimed for consistency, and so chose our data from a single common source: the NCBI RefSeq database (24; release 76, apart from one *Rickettsia peacockii* genome, added from release 77). We began by identifying all eubacterial genera that were represented by multiple named species in RefSeq. We then used Bergey (25), and the wider literature, to classify all species in these genera as “pathogens”, “non-pathogens” or “ambiguous/unknown”. Pathogenicity was defined with respect to vertebrates, not only because vertebrate pathogenicity is better studied, but because vertebrate adaptive immunity is implicated in some theories of genome reduction (7). We note that all such designations must contain an element of uncertainty and ambiguity, not least because of the ubiquity of opportunistic pathogenicity. For this reason, we restricted our definition of “pathogens’’ to species that have repeatedly been reported to cause disease in immuno-competent vertebrate hosts, and preferred species where there was evidence of long-term persistence as a pathogen. For example, we scored *Staphylococcus aureus* as “ambiguous”, because human infection commonly occurs from carriage forms (26), and because carriage status could not be inferred from metadata associated with the sequenced isolates. Similarly, “non-pathogens’’ were defined as species that were known to be free-living or commensals, even if there were isolated cases of secondary infections. For example, *Aeromonas media* was designated as a non-pathogen, despite a single case of isolation, together with pathogenic *Yersinia enterocolitica,* and in a patient recovering from infection with *Aeromonas caviae* (27).

After these assignments, we aimed to choose pairs without further subjectivity, or influence of prior knowledge. As such, we used the following process. First, for each genus containing at least one pathogen and non-pathogen, we downloaded all available genomes (see Table S2). We then used *Phylosift* (28) to align 37 single copy orthologs identified as universal to all bacteria. Concatenated alignments of these loci were checked and corrected by eye, and we used *MEGA7* (29) to build neighbour-joining phylogenies using variation at synonymous sites. We used a modified Nei-Gojobori method using Jukes-Cantor and complete deletion of sites with missing data. These genus-level phylogenies were then midpoint rooted using the *R* package *Phangorn* (30). Then, using these trees, we identified all possible phylogenetically independent pairs of a pathogen and non-pathogen species. This included checking that the genomes from both species were monophyletic with respect to each other, and all other species in the genus-level data set. When a pair included multiple pathogenic or non-pathogenic species, e.g., when a non-pathogen was a sister group to multiple pathogenic species, we chose the best-sampled species with the largest number of available genomes. This process yielded the 31 pairs listed in Table 1 and Table S1. We next noted a suitable outgroup for each pair, and re-estimated trees including only genomes from the pair and outgroup. These pair-level phylogenies were checked for consistency against the relevant whole-genus phylogenies, and used when calculating the independent contrasts (see below).

Finally, we returned to the literature, to identify a subset of pairs with a qualitative difference in ecology between the pathogen and non-pathogen. In particular, we noted pairs where the non-pathogen was extracellular and the pathogen facultatively intracellular (pairs 2, 7, 9, 10, 12 and 23) and where the pathogen, but not the non-pathogen, replicated exclusively within their hosts in nature (pairs 3, 8, 10, 17, 23, 24, 27 and 29); these pairs are indicated in Table 1.

For our *S. suis* data, we used isolates originating from six collections spanning six countries (Table 2) with the same diagnostic criteria of pathogenicity status. The first collection includes isolates from the UK sampled between 2009 and 2011 (described in 5). The second includes isolates from pigs and human meningitis patients from Vietnam, sampled between 2000 and 2010 (described in 5). The third includes carriage isolates from pigs from five intensive farms in the UK, and from five intensive farms and five traditional farms in China from between 2013 and 2014 (described in 31). The fourth includes disease isolates sampled from UK pigs (described in 32). The fifth includes isolates from North American and Spanish pigs, sampled between 1983 and 2016 (33). The sixth includes 29 reference isolates downloaded from GenBank (34). Full details of all genomes are in Table S3.

Pathogenicity status was defined in the following way. Isolates were classified as associated with “disease”, if they were recovered from systemic sites in pigs or humans with clinical signs consistent with *S. suis* infection, including meningitis, septicaemia and arthritis (“systemic disease” isolates), or were recovered from a pig’s lungs in the presence of lesions of pneumonia (“respiratory disease” isolates). Isolates recovered from the tonsils or tracheo-bronchus of healthy pigs or pigs without any typical signs of *S. suis* infection were classified as “carriage”. The remaining isolates remained unclassified, due to insufficient clinical information or ambiguity in the cause of disease.

For most of the collection, serotyping was performed using antisera to known *S. suis* serotypes by the Lancefield method (5, 32). Isolates that could not be typed with known sera were classified as non-typeable. A subset of UK isolates and the Chinese isolates were serotyped *in silico* using capsule genes of known serotypes (described in 31). Isolates that were not serotyped were excluded from comparisons.

Sequence data from all isolates was used to generate *de novo* assemblies using *Spades* v.3.10.1 (35), after first removing low quality reads using *Sickle* v1.33 (36). Measures were taken to ensure all assemblies were high-quality, as described in previous studies (5,31–33). Briefly, Illumina reads were mapped back to the *de novo* assembly to investigate polymorphic reads in the samples (indicative of mixed cultures) using *BWA* v.0.7.16a (37), and genomes that exhibited poor sequencing quality (i.e. poor assembly as indicated by a large number of contigs, low N50 values or a high number of polymorphic reads) or that which were inconsistent with an *S. suis* species assignment were excluded from the analysis. Altogether this left 1,079 genomes.

To identify genetic structure in our *S. suis* isolates we identified 429 low-diversity core genes in our data set, aligned them using *DECIPHER* (38), and stripped regions that could not be aligned unambiguously due to high divergence, indels or missing data. This conserved region of the core genome was first used to construct a consensus neighbour-joining tree using the ape package in *R* and a K80 model (39). This tree is shown in Figures 1b and S1, and was used to generate the covariance matrices used in the phylogenetically corrected regressions (Table S6, Figure S11). The same data were used to identify genetic clusters, using the *hierBAPS* package in *R* (18). Initial analysis identified 35 clusters. To evaluate this clustering we mapped the clusters onto the core gene phylogeny, and following the definition in (40), estimated *F_ST_* between clusters from pairwise nucleotide distances in the core gene alignment. We identified a pair of clusters with very low *F_ST_* (<0.02) that were also monophyletic in the tree, and these clusters were combined to form cluster 3 (Table S1). Full details of all clusters are found in Table S4.

The illustrative genealogy shown in Figure 1c involved mapping to the reference genome BM407 (see Table S3) using *Bowtie2* (41), recombination-stripping in *Gubbins* (41,42), and tree construction with *MrBayes* (43) with default parameters and the HKY+Γ substitution model.

### Genome annotation

For between-species data, we used the *RefSeq* annotations. We also carried out re-annotations using *Prokka* (v2.8.2) (44). While *Prokka* and *RefSeq* annotations were generally congruent, *Prokka* does not explicitly annotate pseudogenes and thus high levels of pseudogenisation in a handful of species (e.g., *Rickettsia prowazekii),* led to erratic results, so we preferred *RefSeq* annotations. We also excluded plasmids because these can be lost during culture and sequencing. However, the main results concerning genome size are robust to their inclusion (Table S5).

The draft *S. suis* genomes were also annotated using *Prokka* (v2.8.2) (44). Orthologous genes were initially identified using *Roary* (45), with the recommended parameter values. We then manually curated these orthology groups, in order to identify orthologous genes that had been wrongly placed in distinct orthology groups either due to high levels of divergence or incomplete assemblies. We also checked all instances of gene absence in each orthology group, since these might have resulted from incomplete genome assemblies. This was undertaken using all-against-all gene group nucleotide *BLAST* search *(BLASTN),* and *BLASTN* search of all orthology groups against all of the genomes in which that group was described as absent (46). The final set of orthology groups were used to define the core genome (Figures S7, S8).

In Figure 3 and related analysis, we defined the “coding” proportion of the genome as any region annotated as a protein- or RNA-coding locus. For both data sets, all genes were assigned a COG category (47), and categories C, E, F, G, H, I, P and Q defined as “metabolic genes”. For the *S. suis* data set we also identified genes that were annotated as transposases or integrases in the *Prokka* annotations.

### Statistical analyses

For the between-species data, each comparison pair differed in the number of genomes sampled, and the amount of evolutionary change between the species. For this reason, we standardized the weightings using a method of independent contrasts. In brief, each comparison point was equivalent to the difference between the ancestral trait values for the sampled genomes from the pathogenic and non-pathogenic species that would be inferred from a Brownian motion model of trait evolution (and using the tip value in the case of a single genome). The contrast for each pair was then standardised by its associated standard deviation. The method used the pic and ace functions in the *ape* package in *R* (39,48), and for each pair, we used the genealogies constructed from the 37 “universal genes”, described above, so that amounts of molecular evolution were comparable across the entire data set. We also added a fixed constant of 1/length(alignment) to deal with zero-length branches in some of the genealogies (reducing this constant by a factor of 10 had no appreciable effect on our results). For each variable, we then tested the validity of the Brownian motion model following the recommendations of (49), and as shown in Figures S4 and S6. For most traits, the model provided a good fit after appropriate transformations (logarithmic for genome size and gene number, and logit for proportions). The sole exception was the proportion of genomes with coding function (Figure S6d-f), which is consistent with the rapid loss of non-functional elements.

Even after standardising the variances, the set of contrasts was usually highly non-normal (e.g., Figure 2a), and so we tested the null hypothesis of a vanishing mean (i.e., no consistent trait difference between pathogenic and non-pathogenic species) by randomly permuting the labels “pathogen” and “non-pathogen” within each pair (i.e., randomly choosing the sign of each of the 31 contrasts). The test statistic was the mean absolute contrast value (using the true signs), and 10^6^ random permutations were used to construct its null distribution. We also repeated results after removing outliers, identified by eye (Table S5). The same permutation approach was used for the within-cluster data set, although here, the disease and carriage isolates were interspersed in the genealogy, and so we used the raw means of the trait values for each class of isolate.

For the between-cluster analyses, tests also had to account for the differences in cluster size. For this reason, most results used weighted linear regression (using the square root of the number of isolates in each cluster as weights). Because these analyses ignored possible covariances between clusters, due to their shared ancestry, we also used phylogenetically-corrected regressions, retaining only the larger clusters (containing at least 20 isolates). These analyses, shown in Table S6 dataset “D”, used the *gls* function in the *nlme* package in *R* (50), and Pagel’s “lambda correlation structure” (51) *(corPagel* in the *ape* package, 39).

## Acknowledgements

We wish to acknowledge Marta Matuszewska and Mukarram Hossain for helpful discussions. LAW, GGRM and ELM were supported by a Sir Henry Dale Fellowship jointly funded by the Wellcome Trust and the Royal Society (109385/Z/15/Z). GGRM was also supported by a ZELS BBSRC award (BB/L018934/1) and a Research Fellowship at Newnham College. JC was supported by an EBPOD fellowship, jointly funded by EBI and the University of Cambridge.

