## Supplementary Figures and Tables for "Genome reduction is associated with bacterial pathogenicity across different scales of temporal and ecological divergence"

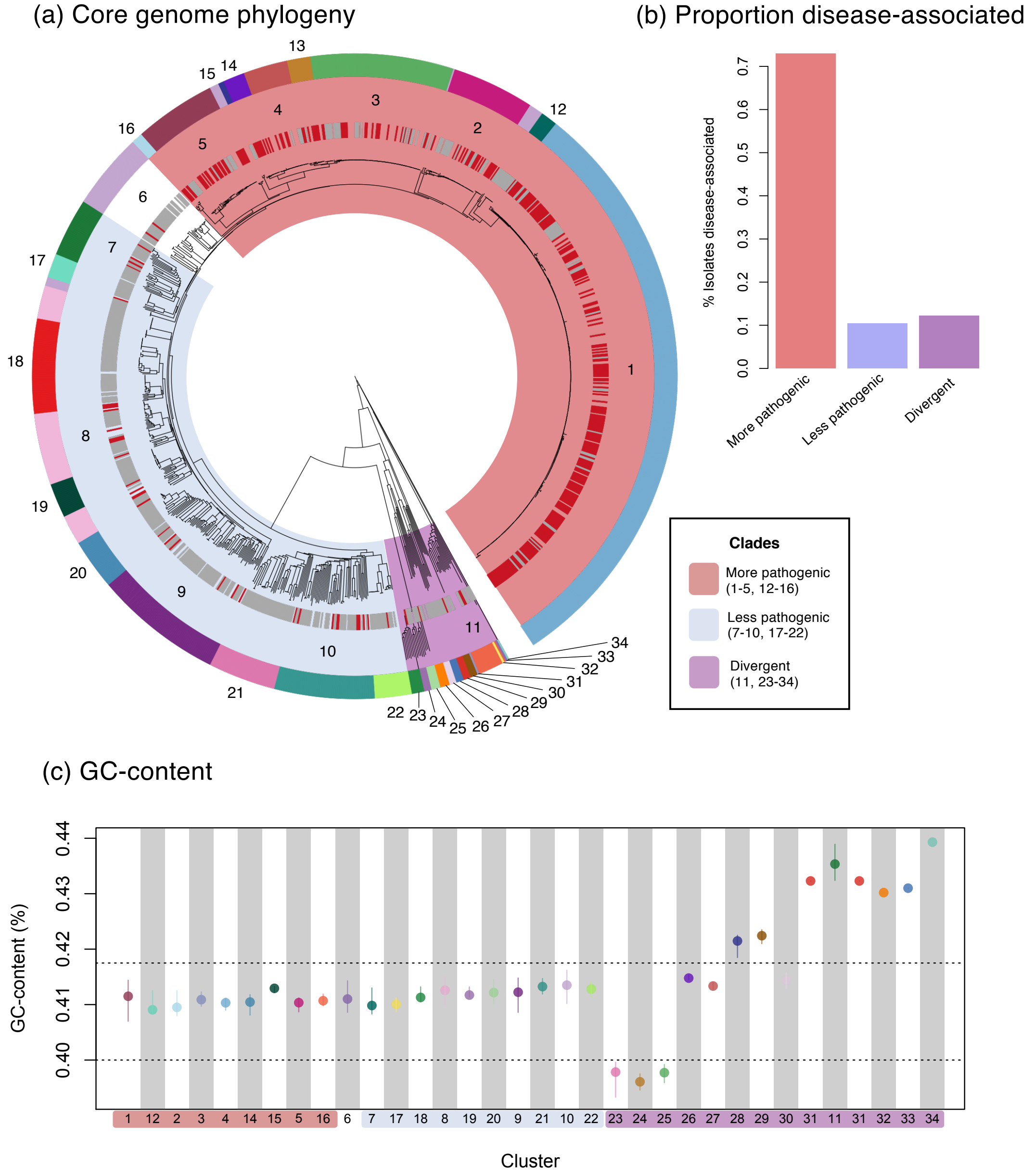


**Figure S1. The genetic structure of *S. suis*.** (a) A core genome phylogeny of our 1079 isolates of *S. suis*. Individual disease (red) and carriage (grey) isolates are indicated in the inner strip. The outermost strip describes the 34 genetic clusters. The 11 "mixed clusters" that include multiple disease and carriage isolates are numbered within the strip (clusters 1-11), and the other 23 clusters are numbered outside the strip (clusters 12-34). The 'more pathogenic' clade is highlighted in red, the 'less pathogenic' clade in blue, and the 'divergent' clade in purple. (b) The proportion of isolates in each clade that are associated with disease (not including isolates for which disease-association is unknown). (c) The whole-genome GC-content for each cluster, with the three clades highlighted. Dashed lines indicate the range of GC values that was used to exclude 10 outlying clusters (3 with low GC, and 7 with high GC), in Figures S7b and S8.

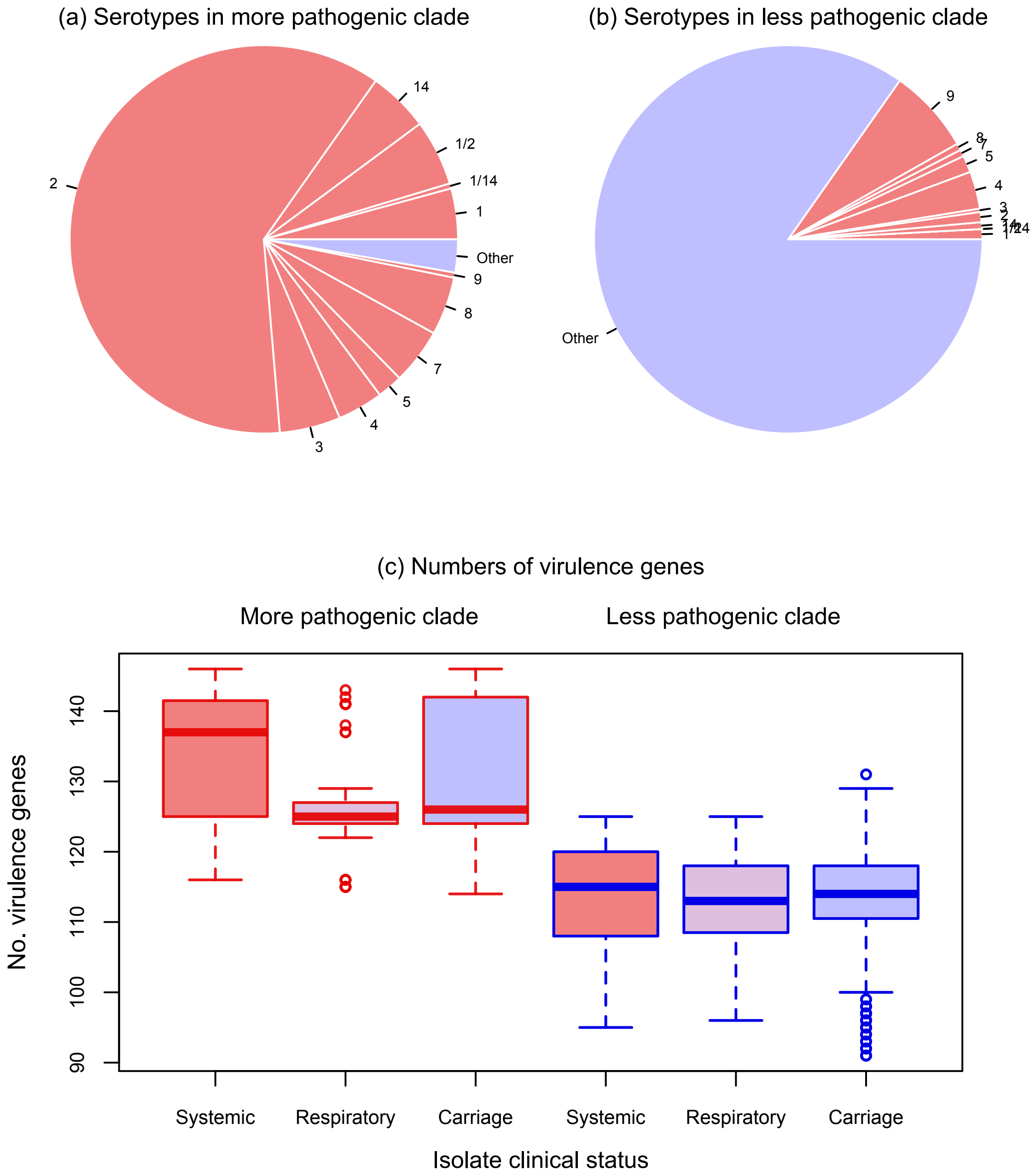


**Figure S2. The reliability of disease-associated phenotyping in the *S. suis* data.** (a) Pie charts showing the proportion of disease-associated serotypes (serotypes 2, 1/2, 1, 14, 3, 4, 5, 7, 8, 9) (53, 54) present in our more and less pathogenic clades. (b) Box plots showing the numbers of virulence genes present in isolates from our more and less pathogenic clades, and in isolates with different clinical statuses within these clades. This is based on a set of 157 virulence genes that were identified in several previous studies of *S. suis* virulence (53, 55–58).

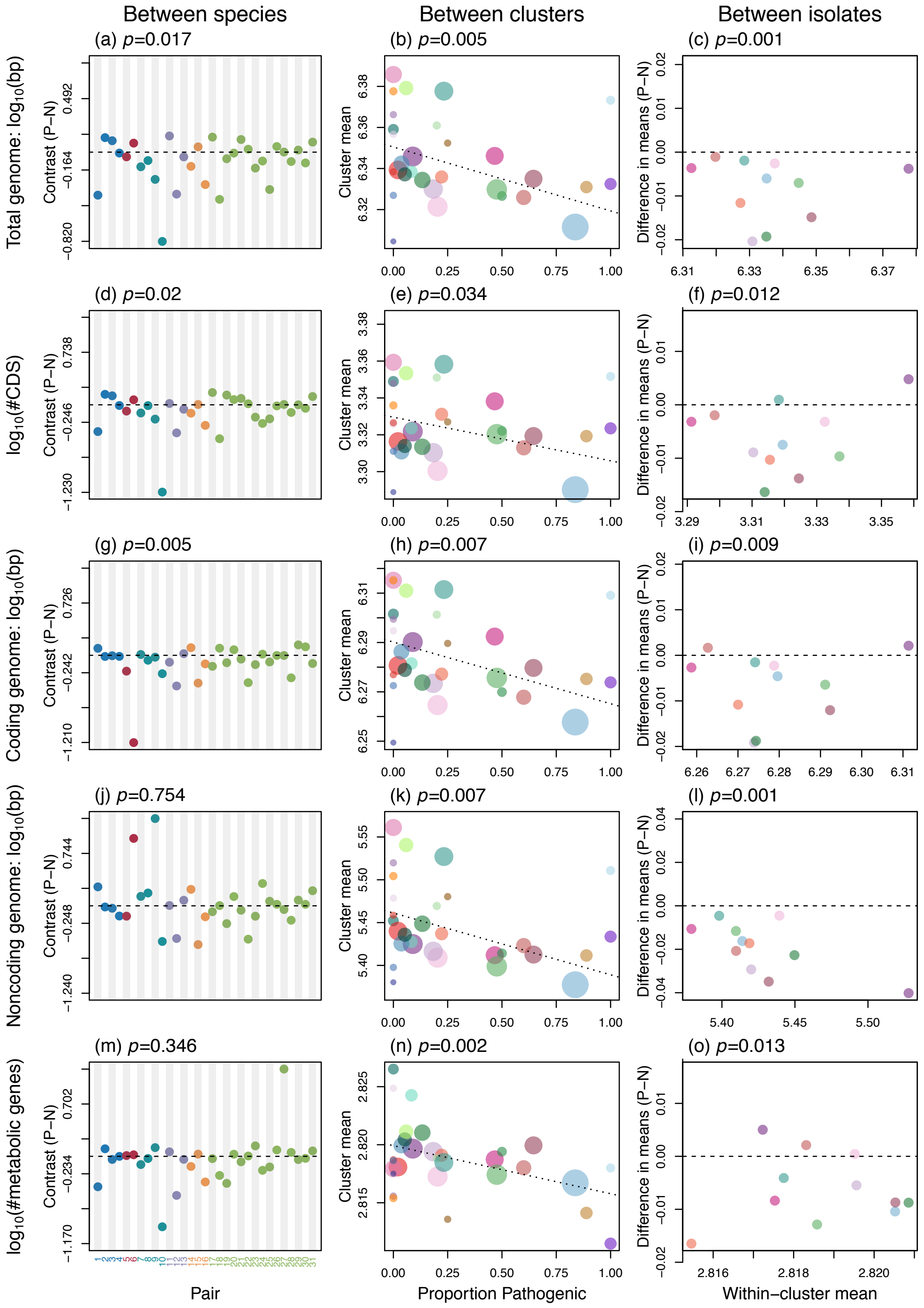


**Figure S3. The association of pathogenicity with different measures of genome size (a-i), and the endosymbiont syndrome (j-o).** (a-c) Total genome size after including plasmids (note that this makes no difference for the *S. suis* data set); (d-f) Total number of intact protein-coding genes; (g-i) Total length of protein and RNA-coding elements; (j-l): Length of genome non coding for proteins or RNAs; (m-o) The number of intact protein-coding genes with metabolic function. All other details match Figure 2.

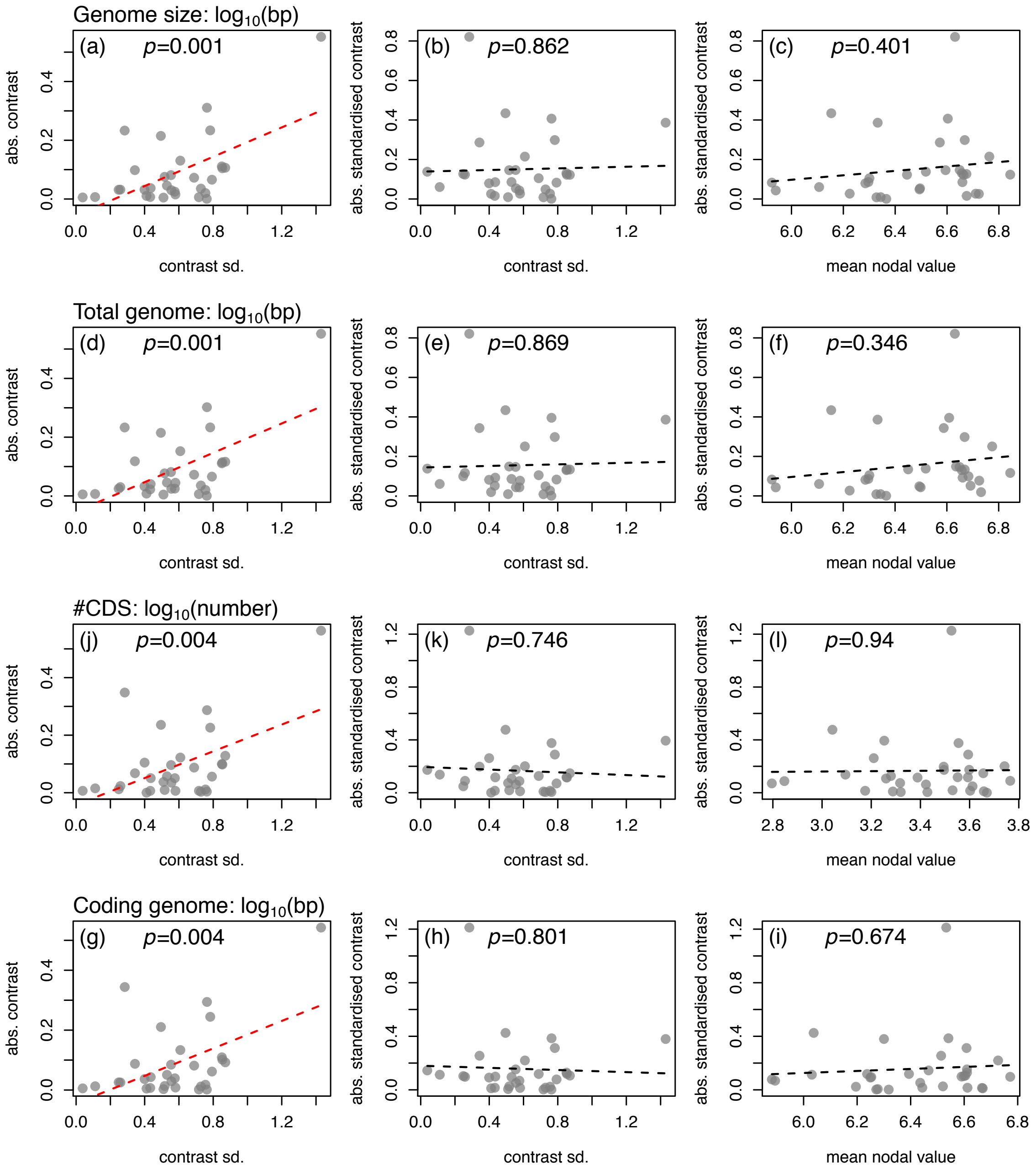


**Figure S4. Adequacy of the Brownian Motion model used to generate between-species independent contrasts for genome size.** The left-hand column plots the absolute values of the raw (unstandardised contrasts) against the contrast standard deviations, as inferred from the trait values and bacterial genealogy. A significant positive regression slope is consistent with the Brownian Motion model. The middle and right-hand columns plot the absolute value of the standardised contrast against its standard deviation, or the mean trait value for the two relevant nodes of the genealogy. The lack of a relationship, in each case, is consistent with the Brownian Motion model, and implies that all points should have equal weight in the tests. Each column corresponds to a different measure of genome size, matching those used in Figure 2 (a-c), or Figure S3 (d-i).

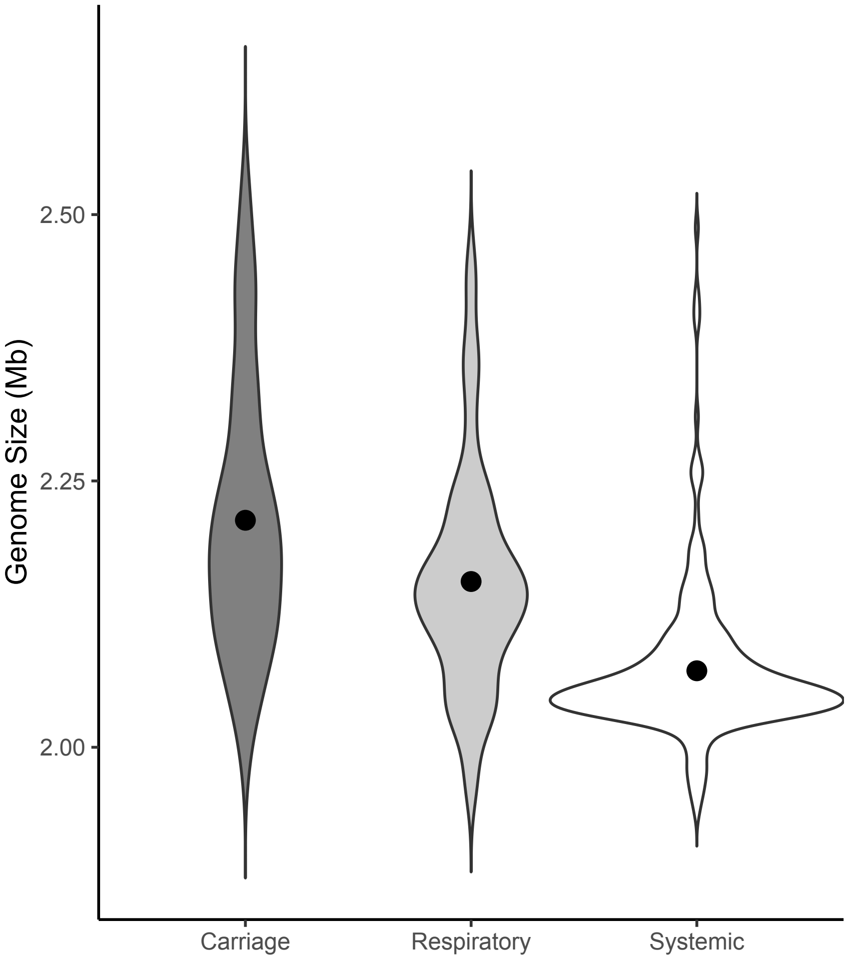


**Figure S5. Genome size and disease severity in *S. suis.*** Violin plots of the distribution of genome sizes for the three classes of *S. suis* isolates used in this study. Across the data set as a whole, there is a tendency for the most serious systemic disease isolates to have the smallest genomes, while isolates associated with less serious respiratory disease have genomes of intermediate size.

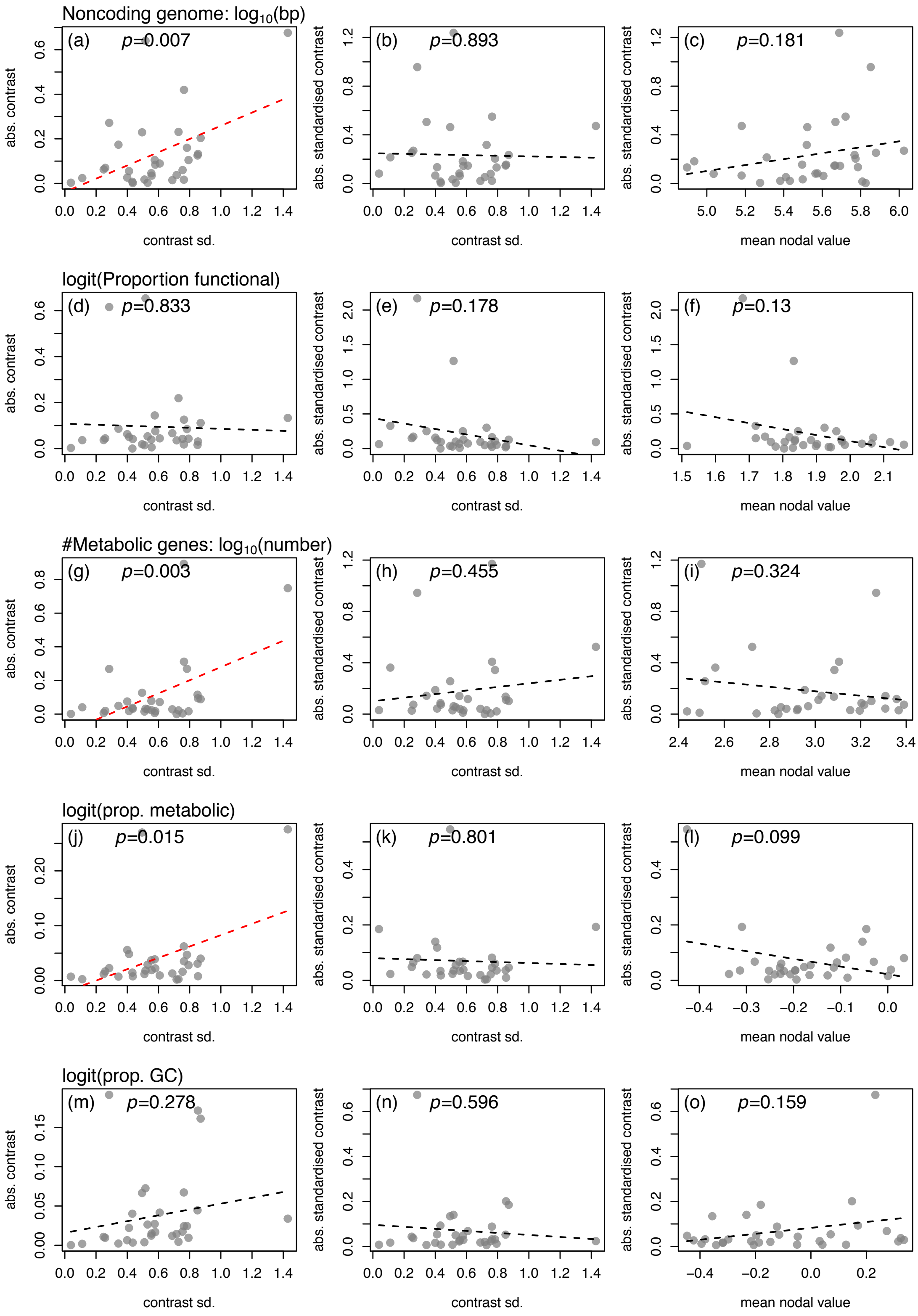


**Figure S6. Adequacy of the Brownian Motion model used to generate between-species independent contrasts for measures of the endosymbiont syndrome.** All details match Figure S4, except for the trait values used, which match those in Figure 3 (d-f, j-o) or Figure S3 (a-c, g-i). Of note is the failure of the Brownian motion model to fit the proportion of the genome that is functional (d-f).

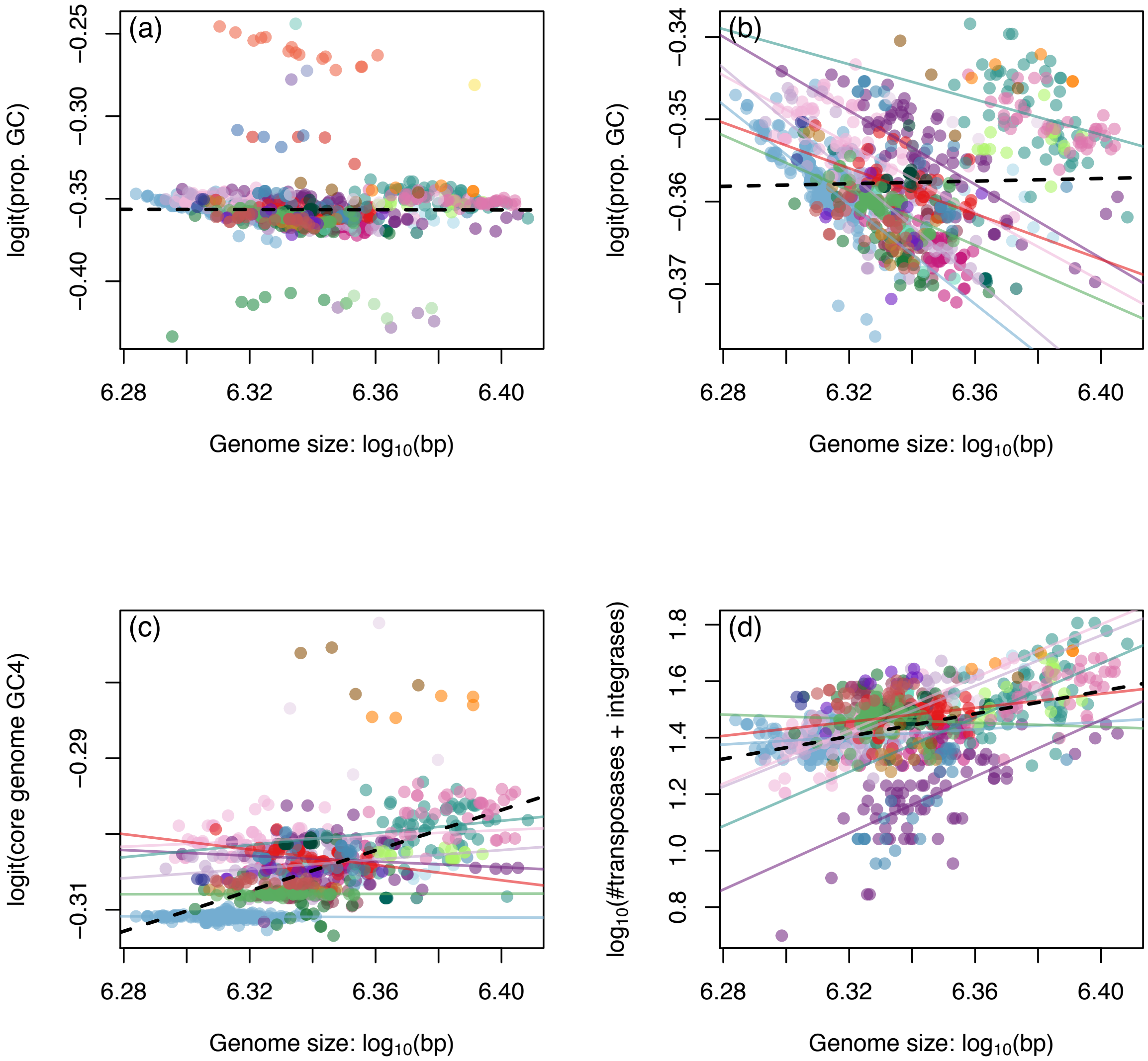


**Figure S7. Variation in genomic GC content within the *S. suis* data. Each point represents a single *S. suis* isolate, coloured according to the genetic clusters from Figure 1b, and all lines represent best-fit regression slopes using ordinary least-squares.** (a) Interpreting the relationship between GC-content and genome size is complicated by divergent clusters with anomalous GC values (see Figure S1). (b) After removing these clusters, there is not clear association in the data as a whole (dashed line), but equally clear negative trends within each of the clusters (coloured lines show results for the 7 largest clusters). (c) These patterns change when we consider only sites in the core genome (i.e., sites present in all isolates). In particular, for the most rapidly evolving fourfold degenerate sites in protein coding genes, there is little variation in GC4 within clusters (coloured solid lines), but a positive relationship across the data as whole (black dashed line). (d) These patterns are explained by the rapid dynamics of mobile elements, with transposes and integrases more common in the larger genomes. The differential presence of these GC-poor elements counteracts the trend seen in the core genome.

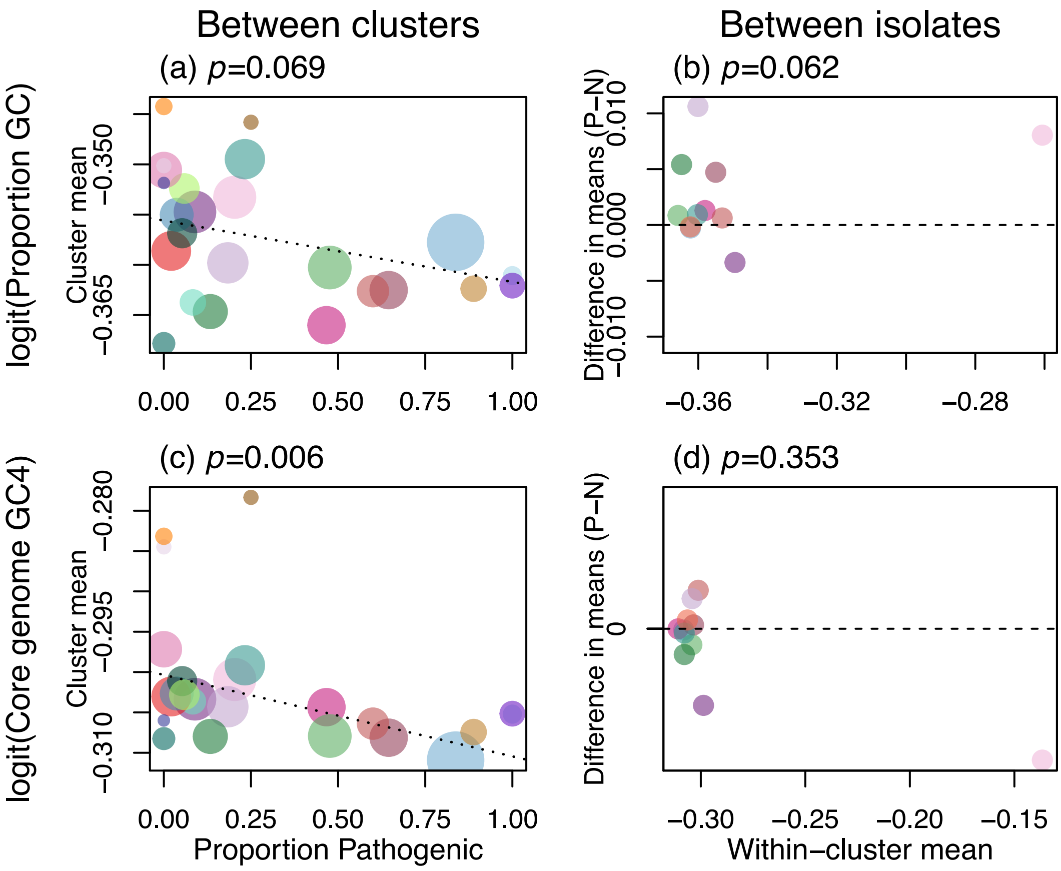


**Figure S8. GC content and pathogenicity *in S. suis*, removing the divergent clusters.** Results are shown for *S. suis* after the removal of 10 divergent clusters with anomalous GC contents (see Figure S1). (a-b) overall genomic GC content, as in Figure 3h-i; (c-d) GC content at four-fold degenerate sites in protein-coding genes from the core genomes (i.e., that are present in all isolates). Core genome sites show the predicted association of low-GC with pathogenicity, but the effect is only evident over long periods of time (c). In the genome as a whole, the effect is masked by the absence of GC-poor mobile elements in smaller genomes.

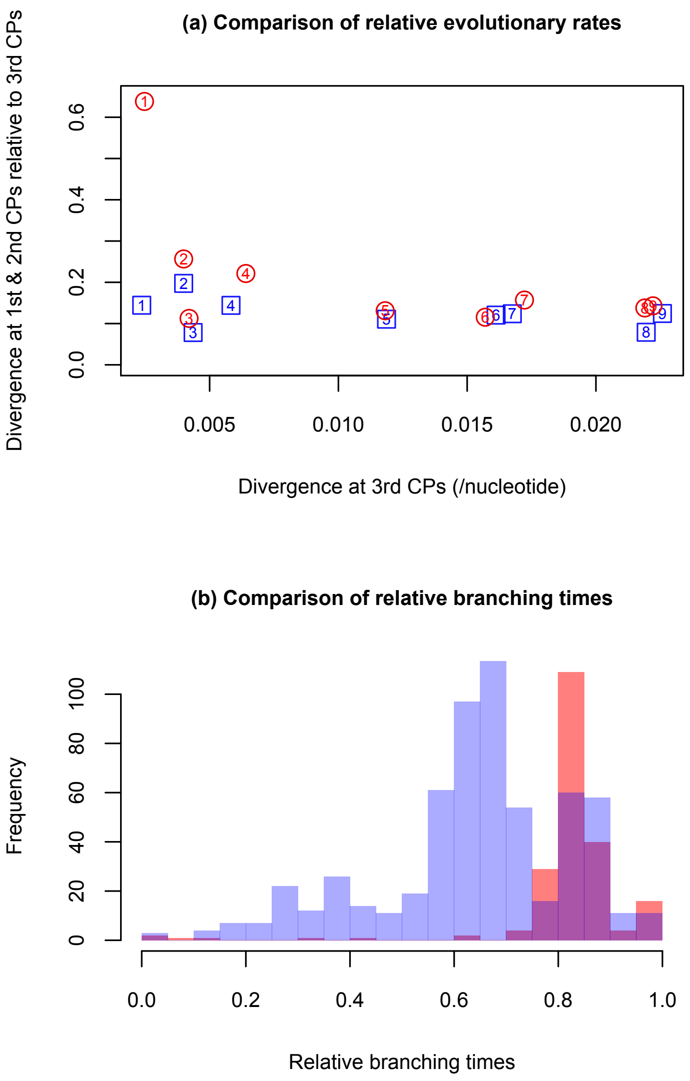


**Figure S9. Evidence of ineffective purifying selection in pathogenic *S. suis*.** (a) Comparison of evolutionary divergence at 1st and 2nd codon positions relative to divergence at 3rd codon positions in 9 distance-matched clusters from more (red circles) and less (blue squares) pathogenic clades. As the phylogenetic relationships between isolates are difficult to reconstruct accurately due to high rates of recombination both within and between populations (5), we compared the divergence of single pairs of isolates from each cluster. Estimates from each cluster from the pathogenic clade were taken from the most divergent pair of isolates within that cluster, and compared with a pair of isolates from a cluster from the less pathogenic clade that were the closest match for distance at 3rd codon positions. In 8/9 matched comparisons the cluster from the more pathogenic clade has the greater divergence at 1st and 2nd codon positions relative to divergence at 3rd codon positions. (b) The distribution of average branching times for our more (red) and less (blue) pathogenic clusters as estimated by the *ltt* function in the *phytools* package in *R* (59) applied to a midpoint-rooted neighbour-joining tree of our core gene alignment. A non-parametric rank test shows significant difference between the two groups (Wilcoxon rank test, $p = 2.5 \times 10^{-8}$). This difference is robust to the inclusion/exclusion of disease-associated isolates.

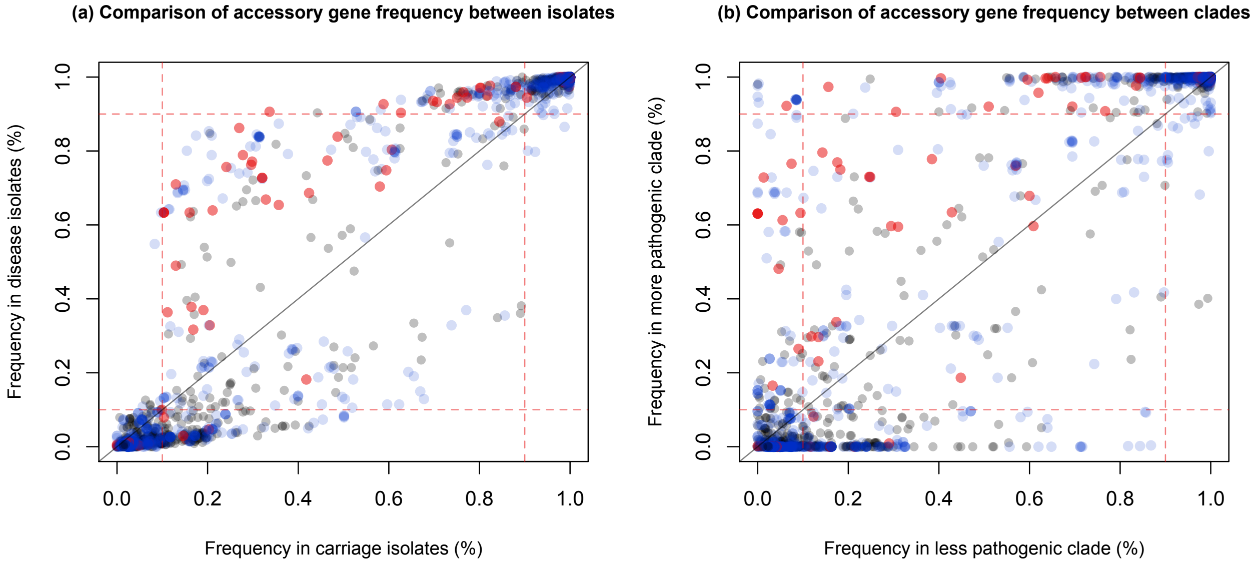


**Figure S10. Patterns of gene presence/absence in *S. suis*.** Comparisons of the frequency of homologous genes across (a) all disease-associated isolates and all carriage isolates, and (b) across the more and less pathogenic clades. Solid lines indicate equal frequencies, and red dashed lines indicate frequencies of 10% and 90% in either category. Each point represents a homologous gene. Metabolism genes (identified through our COG classification) are shown in blue, virulence genes (identified in previous studies) in red, and all other genes in grey. Gene frequencies are strongly correlated across carriage and disease isolates. No genes that are common in carriage isolates are absent from disease isolates (or vice versa). While gene frequencies are more divergent across the more and less pathogenic clades, no genes that are present in >90% of isolates in the less pathogenic clade are uniformly absent from isolates in the more pathogenic clade. This suggests that genome reduction is largely driven by the loss of broadly non-essential genes.

**Table S1.** Details of 31 phylogenetically-independent species pairs.

**Table S2.** Details of 1,318 genomes used in the between-species analysis.

**Table S3.** Details of 1,079 *S. suis* genomes.

**Table S4.** Details of *S. suis* genetic clusters.

**Table S5.** Robustness of between-species analyses.

**Table S6.** Robustness of the association between genome size and pathogenicity in between-cluster data.
