## Supplementary Table 4 for "Genome reduction is associated with bacterial pathogenicity across different scales of temporal and ecological divergence"

**Table S4.** Details of *S. suis* genetic clusters

| # | Genome size range (Mb) | # Isolate: Total (SP, RP, N) | Origin(s) | Dates |
|---|---|---|---|---|
| 1 | 1.92 - 2.19 | 328 (199, 12, 41) | VN, UK, CA, CN, ES | 2000 - 2016 |
| 2 | 2.09 - 2.29 | 45 (2, 12, 16) | UK, CA | 2010 - 2016 |
| 3 | 2.04 - 2.26 | 78 (2, 18, 22) | UK, CA, US, CN | 1988 - 2016 |
| 4 | 2.04 - 2.19 | 24 (4, 2, 4) | UK, ES | 2010 - 2016 |
| 5 | 2.10 - 2.26 | 45 (11, 9, 11) | CA, UK, US, CN | 1990 - 2016 |
| 6 | 1.96 - 2.30 | 61 (6, 3, 40) | UK, CN, CA | 1987 - 2016 |
| 7 | 2.01 - 2.31 | 32 (2, 2, 26) | UK, CN, CA | 2008 - 2016 |
| 8 | 1.98 - 2.36 | 73 (8, 4, 47) | UK, CA, CN, US, ES | 1983 - 2016 |
| 9 | 1.99 - 2.54 | 70 (2, 3, 51) | UK, CA, CN, US | 1989 - 2016 |
| 10 | 2.21 - 2.55 | 55 (3, 7, 33) | UK, CA, CN, US | 1988 - 2016 |
| 11 | 2.04 - 2.29 | 15 (1, 1, 7) | UK, CN, CA | 2009 - 2014 |
| 12 | 2.07 - 2.36 | 9 (0, 0, 9) | CN, CA | 2013 - 2016 |
| 13 | 2.05 - 2.30 | 13 (7, 1, 1) | UK, CA | 2010 - 2013 |
| 14 | 2.05 - 2.30 | 12 (4, 4, 0) | CA, UK | 1989 - 2016 |
| 15 | 2.01 - 2.02 | 3 (0, 0, 2) | CA | 1991 - 2016 |
| 16 | 2.27 - 2.42 | 6 (1, 3, 0) | CA, UK | 1989 - 2014 |
| 17 | 2.13 - 2.26 | 13 (0, 1, 11) | UK, CA | 1994 - 2016 |
| 18 | 2.04 - 2.21 | 52 (0, 1, 46) | CN, UK, CA | 2009 - 2013 |
| 19 | 2.14 - 2.21 | 19 (0, 1, 18) | CN | 2013 |
| 20 | 2.10 - 2.45 | 28 (0, 1, 26) | CN, CA, UK | 1989 - 2016 |
| 21 | 2.28 - 2.53 | 37 (0, 0, 37) | CN | 2013 - 2014 |
| 22 | 2.30 - 2.48 | 20 (1, 0, 16) | CN, UK, ES, CA | 2011 - 2016 |
| 23 | 1.97 - 2.24 | 7 (1, 0, 1) | UK, CA | 2008 - 2014 |
| 24 | 2.23 - 2.39 | 4 (0, 0, 3) | UK, CA | 2013 - 2014 |
| 25 | 2.24 - 2.39 | 5 (1, 0, 4) | UK, CA | 2010 - 2016 |
| 26 | 2.29 - 2.46 | 5 (0, 0, 3) | CA, US | 2016 |
| 27 | 2.15 - 2.40 | 4 (0, 0, 4) | UK, CN | 2013 |
| 28 | 2.09 - 2.26 | 4 (0, 0, 4) | CN, UK | 2013 - 2014 |
| 29 | 2.07 - 2.17 | 4 (0, 0, 4) | CN, CA | 2013 - 2016 |
| 30 | 2.17 - 2.36 | 4 (1, 0, 3) | CA, UK, CN | 1988 - 2013 |
| 31 | 2.18 | 1 (0, 0, 0) | UK | 2010 |
| 32 | 2.46 | 1 (0, 0, 1) | CN | 2013 |
| 33 | 2.15 | 1 (0, 0, 1) | UK | 2013 |
| 34 | 2.16 | 1 (0, 0, 1) | CN | 2013 |

**Note:** SP: Systemic pathogen, RP: Respiratory pathogen, C: Carriage; VN: Vietnam, UK: United Kingdom, CA: Canada, CN: China, ES: Spain.
