## Supplementary Tables 5 & 6 for "Genome reduction is associated with bacterial pathogenicity across different scales of temporal and ecological divergence"

**Table S5: Robustness of between-species analyses**

| Trait | Outlier(s) removed $p$ [$n$] | "changed ecology" removed $p$ ($n = 19$) | "changed ecology" only $p$ ($n = 12$) |
|---|---|---|---|
| Genome size, $\log_{10}(bp)$ | 0.0307* [30] | 0.0180* | 0.4526 |
| Total genome size (incl. plasmids) $\log_{10}(bp)$ | 0.0321* [30] | 0.0264* | 0.3774 |
| Coding genome size, $\log_{10}(bp)$ | 0.0102* [30] | 0.0210* | 0.1300 |
| #CDS, $\log_{10}(number)$ | 0.0414* [30] | 0.0480* | 0.2790 |
| #Metabolic genes, $\log_{10}(number)$ | 0.0148* [30] | 0.0212* | 0.8091 |
| logit(proportion metabolic) | 0.0979 [30] | 0.9855 | 0.7803 |
| Non-coding genome size, $\log_{10}(bp)$ | 0.2312 [29] | 0.7728 | 0.5670 |
| logit(proportion functional) | 0.3431 [29] | 0.3095 | 0.2008 |
| logit(proportion GC) | 0.0404* [30] | 0.3087 | 0.0487* |

**Note**: Permutation tests of the difference in trait values between pathogenic and non-pathogenic sister species, calculated using standardised independent contrasts. The first column shows results after the removal of outlying points (identified visually). The second columns show subsets of the data, separating the $n = 12$ points with a changed ecology, defined as pairs where the pathogen species was scored as facultatively intracelluar and the non-pathogen as extracellular, and/or the pathogen as host-restricted, and the non-pathogen as not host restricted. * $p < 0.05$.

**Table S6: Robustness of the association between genome size and pathogenicity in between-cluster data**

| Dataset: #clusters (# genomes): | A: 33 (1078) | B: 24 (883) | C: 31 (1060) | D: 14 (948) |
|---|---|---|---|---|
| | $\hat{\beta}$ ($p$) | $\hat{\beta}$ ($p$) | $\hat{\beta}$ ($p$) | $\hat{\beta}$ ($p$) [$\hat{\lambda}$]; GLS regression |
| Genome size, $\log_{10}(bp)$ | -0.031 (0.0054) | -0.042 (0.0022) | -0.042 (0.0008) | -0.032 (0.0000) [1.16] |
| Coding genome size, $\log_{10}(bp)$ | -0.025 (0.0067) | -0.034 (0.0020) | -0.034 (0.0013) | -0.043 (0.0086) [1.14] |
| #CDS, $\log_{10}(number)$ | -0.024 (0.0341) | -0.038 (0.0060) | -0.035 (0.0044) | -0.049 (0.0034) [1.15] |

**Note**: A: Dataset contains the 33/34 genetic clusters containing isolates that could be unambiguously assigned as "clinical" or "non-clinical"; B: Dataset retains only clusters for which at least 2/3 of isolates could be unambiguously assigned; C: Measures of genome size were calculated exclusively from isolates assigned as "non-clinical" (which were found in 31/34 clusters); D: Dataset contains only "large" clusters, which contained at least 20 isolates. All entries report results from linear regression of mean trait value onto the proportion of clinical strains in each cluster, using sample-size-weighted least squares (datasets A-C), or generalized-least squares, correcting for covariation due to shared ancestry (dataset D). $\hat{\beta}$: best-fit regression slope; $p$: the associated $p$-value; $\hat{\lambda}$: the best-fit value of Pagel's $\lambda$ for phylogenetically-corrected regression.
